# The Muscarinic Acetylcholine Receptor in Dermal Papilla Cells Regulates Hair Growth

**DOI:** 10.1101/2024.04.26.591334

**Authors:** Gary K.W. Yuen, K.W. Leung, Queenie W.S. Lai, Maggie S.S. Guo, Alex X. Gao, Janet Y.M. Ho, Harry C.T. Chu, Karl W.K. Tsim

## Abstract

The role of cholinergic system in hair biology is poorly understood. In M4 muscarinic receptor (mAChR) knockout mice, the hair follicles have a prolonged telogen phase and fail to produce hair shafts. Here, we reported that hair growth was regulated by cholinergic system via mAChRs. Dermal papilla cells expressed different cholinergic biomarkers. Inhibiting AChE or activating mAChR in dermal papilla cells, culture vibrissae and skin epidermis promoted the hair growth. In cultured papilla cells treated with bethanechol, an agonist of mAChR, an activation of Wnt/β-catenin signalling was illustrated by various indicative biomarkers, including phosphorylation of GSK-3β and mRNA expression of various molecules for Wnt/β- catenin signalling. Activation of Wnt/β-catenin signalling was mediated by PI3K/AKT and ERK signalling upon the stimulation of bethanechol. In addition, an increase in hair shaft elongation was observed in mouse vibrissae upon the treatment of bethanechol, suggesting the cholinergic role in hair growth.

## INTRODUCTION

In non-neuronal cells, the cholinergic system, composing acetylcholine (ACh) synthesizing enzymes, transporters, receptors, and degrading enzymes, exhibits a wide range of biological roles on different organs and pathologies (Beckmann & Lips, 2013; Grando, 2007; Kurzen et al., 2007; Wessler & Kirkpatrick, 2008). Several lines of evidence suggest that the dysregulation of cholinergic system in different tissues is leading to diverse diseases, having significant implications for human health. For an example in skin epidermis, the role of cholinergic molecules has been proposed to be involved in different skin functions (Grando et al, 1993; Guo et al, 2023). A synapse-like interplay between melanocyte and keratinocyte, named as “skin synapse”, has been proposed (Wu et al, 2020a; Wu et al, 2018; Wu et al, 2020b). The direct involvement of cholinergic molecules is to regulate skin melanogenesis in both melanocyte and keratinocyte. Under the exposure to UV, keratinocyte is stimulated to release ACh acting on acetylcholine receptor (AChR) localized on melanocyte to trigger the synthesis and release of melanin. In parallel, the patients suffering from atopic dermatitis showed over 14-fold higher levels of ACh in epidermis and dermis, as compared to normal levels (Heyer et al, 1997; Wessler et al, 2003).

In mammalian skin, hair follicle is an organ found in skin dermal layer. The hair follicle, containing over 20 different cell types, is responsible for hair growth via a complex and dynamic process (Bernard, 2006). The hair growth is tightly regulated by the follicle in a cyclic manner (Lin et al, 2022), which consists of three distinct phases, namely: (i) anagen (the active hair growth phase); (ii) catagen (the regression phase); and (iii) telogen (the resting phase), which are characterized by unique morphological and molecular features (Oh et al, 2016; Paus & Cotsarelis, 1999). To achieve a well-balanced between hair growth and regression, a range of growth inhibiting and/or promoting signals are involved in regulating different phases of hair follicle. Among those growth promoting signals, wingless (Wnt), fibroblast growth factor (FGF), vascular endothelial growth factor (VEGF), and insulin-like growth factor 1 (IGF-1), have been identified as key players in promoting hair growth (Lin et al, 2015; Natarelli et al, 2023; Yano et al, 2001). In contrast, the inhibiting signals, such as transforming growth factor β (TGF-β) and bone morphogenetic protein (BMP), are known to drive hair follicles into regression (Hibino & Nishiyama, 2004; Rendl et al, 2008; Sharov et al, 2005). Activation of Wnt/β-catenin signaling triggers the growth of hair follicles by stimulating different gene expressions at the anagen phase (Andl et al, 2002; Beaudoin III et al, 2005; Huelsken et al, 2001; Kishimoto et al, 2000). The dermal papilla cell (DPC), a group of specialized mesenchymal cells clustering at the base of hair follicle, is a crucial player in regulating hair growth (Yang & Cotsarelis, 2010): this cell secretes a range of paracrine and autocrine factors, including Wnt, R-spondin, FGF, and Noggin, acting on follicular bulge stem cells to initiate the anagen phase, as well as to promote hair growth (Driskell et al, 2011; Matsuzaki & Yoshizato, 1998; Morgan, 2014).

The role of the cholinergic system in hair biology remains an interesting topic with limited finding, having only a few reports exploring the relationship of the complex process. Several lines of evidence support the notion of ACh playing role in hair biology. Alzheimer’s patients taking oral reversible and competitive acetylcholinesterase (AChE) and butyrylcholinesterase (BChE) inhibitor, rivastigmine, exhibited symptoms of hypertrichosis and hair re-pigmentation (Chan et al, 2020; Imbernón-Moya et al, 2016; Müller, 2007). In DPCs treated with AChE inhibitor, nor-galantamine, a stimulation of anagen’s activating signalling has been identified (Kowal et al, 2018; Prvulovic et al, 2010; Scott & Goa, 2000; Yoon et al, 2019). In addition, the knockout mice of M4 muscarinic receptor (M4 mAChR) showed a defect in hair growth: the hair follicles had a significantly prolonged telogen phase, and which failed to produce pigmented hair shafts (Hasse et al, 2007; Takahashi, 2021). These findings provide supporting evidence that the cholinergic system may play roles in the multifaceted processes of hair growth. Here, we hypothesize that the cholinergic signalling in hair follicles is able to regulate hair growth. The expression profiles of cholinergic molecules in DPCs and hair follicles were determined. In addition, the release of ACh, induced by solar light, from DPC was demonstrated, which thereafter activated the AChRs localized on DPCs; the cholinergic activation triggered the Wnt/β-catenin signalling in order to stimulate hair growth. These findings shed light on the role of cholinergic system in hair growth.

## RESULTS

### Characterization of cholinergic molecules in DPC

DPC was employed here as the major cell type in testing the induction of hair growth. The cultures showed a rapid growth from day 2 to 6 after plating: the maximal cell number was at day 7 (**Supplementary Fig. 1a**). DPC cultures contained mainly the enzymatic activity of AChE instead of BChE, as revealed by the effect of different specific inhibitors (i.e., BW284c51 as the AChE inhibitor and iso-OMPA as the BChE inhibitor) (**Supplementary Fig. 1b**). The expressions of cholinergic molecules in cultured DPCs were identified, showing changes during growth. Specifically, the protein expression of AChE decreased, while BChE increased as the culture time progressed (**Fig. 1a**). This indicated a potential regulatory mechanism governing the expressions of these two enzymes.

**Figure 1.**
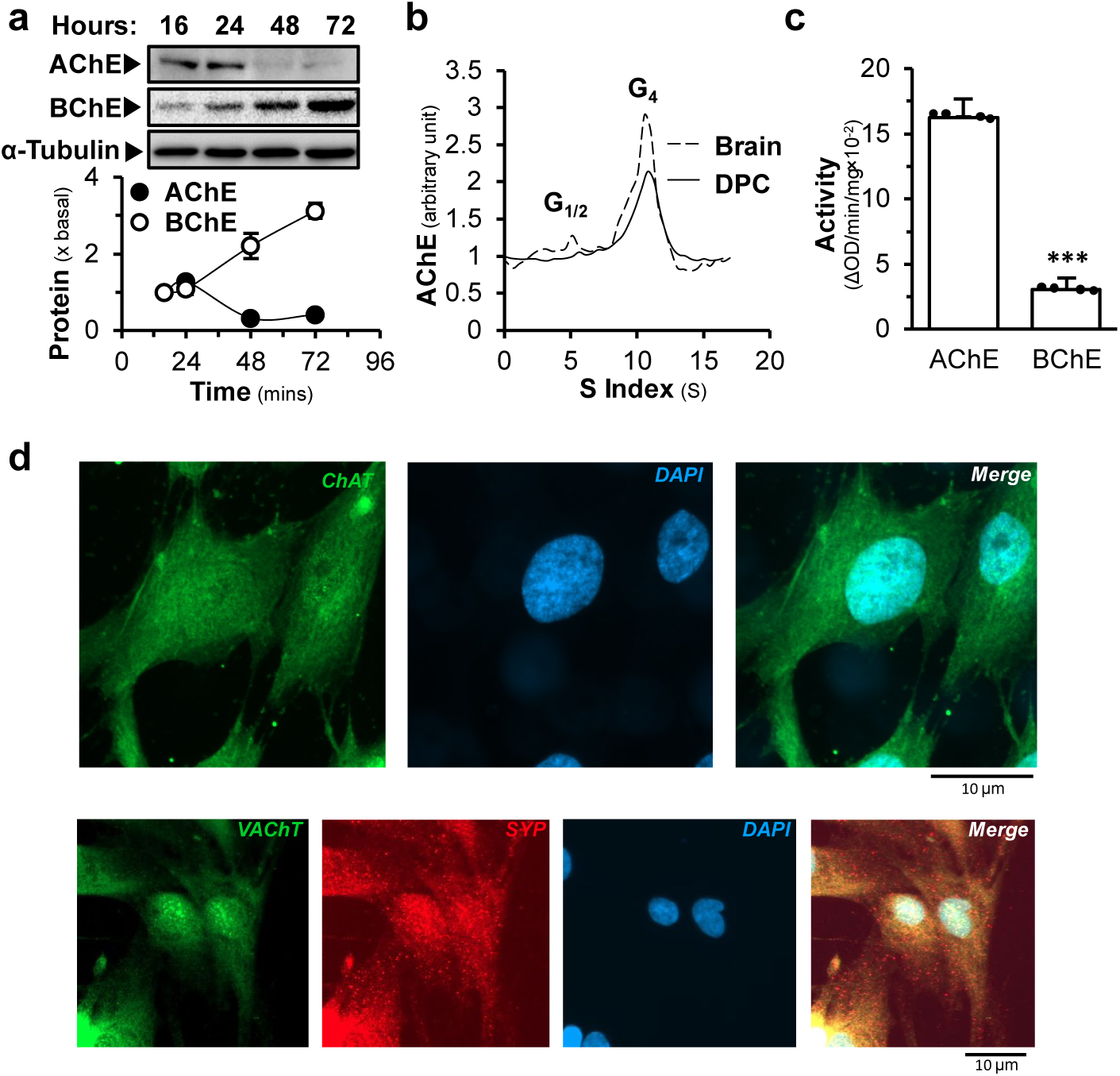
Expression of cholinergic molecules in DPC. Cultured DPCs were harvested on different days, and cell lysate and mRNA were collected. **(a)** cell lysates (20 μg) were subjected to Western blot analysis. AChE (∼70 kDa), BChE (∼ 40 kDa), α-tubulin (∼50 kDa; a loading control) are shown (upper) and quantified (lower) (*n*=3). **(b)** AChE molecular forms in a sucrose density gradient analysis. The sedimentation (S) value was estimated from the position of the sedimentation markers (*n*=3). **(c)** The activities of AChE and BChE in DPC cultures were determined (*n*=4). **(d)** Immunocytochemical staining of ChAT (green), VAChT (green), SYP (red) and DAPI staining (blue) in DPC (*n*=4). Data are normalized and expressed as the % increase, or % of control, or fold of basal (x basal), or in relative amount in comparison to control, in mean ± SEM, * *p* < 0.05, ** *p* < 0.01, *** *p* < 0.001.

In neuronal synapses, globular tetramer (G_4_) form of AChE is composed of four AChE catalytic subunits. With the help of proline-rich membrane anchor, G_4_ form of AChE attaches to cell membrane and hydrolyses ACh, extracellularly. We suspected that AChE in DPC has similar function in hair follicles. AChE form in the DPC culture was determined by a sucrose density gradient, showing the presence of amphiphilic monomer (G_1_), dimer (G_2_) and tetramer (G_4_) (**Fig. 1b**). Notably, the G_4_ AChE was the major form. The mRNA expression of BChE was higher than that of AChE by over 30% (**Supplementary Fig. 1c**). In contrast, the enzymatic activity of AChE was ∼ 6 times higher than that of BChE (**Fig. 1c**). The hydrolysing rate of ACh was much faster by AChE than BChE, and therefore, AChE should still play a dominant role in breaking down ACh here. The mRNA expressions of AChE, BChE and ChAT in different cell lines were compared by qPCR. AChE mRNA expression of DPC was much lower than that of HaCaT and SH-SY5Y cells (**Supplementary Fig. 1d**). In addition, the identification of subunits for nAChR and mAChR, as well as other cholinergic molecules in DPC, was done by specific primers in PCR. **Supplementary Fig. 2a-c** shows the presence of these molecules.

### DPC as a source of ACh

ChAT is an enzyme known for its pivotal role in the synthesis of ACh. While vesicular ACh transporter (VAChT) is a crucial transporter responsible for sequestration of ACh into secretory organelles, i.e., synaptic vesicle, in neurons. To confirm the presence of cholinergic molecules in DPC, immunofluorescent staining was performed. In the present of Triton X-100, the expressions of ChAT, VAChT and synaptophysin (SYP) in DPCs were recognized by using anti-ChAT, anti-SYP and anti-VAChT antibodies (**Fig. 1d**). The staining of SYP was localized intracellularly in a punctate manner. Dermal papilla is the regulating centre in hair follicle, which secretes growth factors to stimulate the development of hair follicle. Here, we hypothesized that DPC could be the source of ACh in activating cholinergic system in hair follicle. To support this hypothesis, the light sensitive protein should be present in DPC. Opsin has been proposed to sense light by skin surface, as to cause ACh release from skin epidermal cells (Wu et al., 2020a). The mRNAs encoding OPN1, OPN2 and OPN3 were identified in DPC, while the expressions of other opsins were not observed. The expression profile of these opsin was peaked at 1 day after culture (**Fig. 2a**). To reveal the possible release of ACh from DPC, KCl was employed to depolarize the cell membrane (Pincus & Weinfeld, 1984). Starting from the concentration of 25 mM KCl, an increase of ACh in the conditioned medium, released by DPC culture, was detected by an increase of ∼2 folds (**Fig. 2b**). The treatment with KCl in DPCs resulted in the release of ACh over time, reaching a maximum at 15 min of the challenge (**Fig. 2c**). In addition, the exposure of solar light to DPCs was able to cause the release of ACh, and which was in a time-dependent manner, having a maximum at 5 min of the exposure (**Fig. 2d**). EDTA, a Ca^2+^ chelator, was able to block the light-mediated release of ACh, suggesting the involvement of Ca^2+^ in the ACh release. In line to this notion, the amount of ACh in P7 anagen was ∼5 times higher than that of the P23 telogen in skin tissues (**Fig. 2e**). Thus, the level of ACh in mouse skin was varied during the hair cycle development. Besides, the pTOPflash plasmid containing TCF-binding sequence for β-catenin TCF complex was used to quantify the activation of Wnt/β-catenin, a key signal during hair growth. The exposure of solar light in pTOPflash-transfected DPC was able to increase the luciferase activity in a time-dependent manner (**Fig. 2f**).

**Figure 2.**
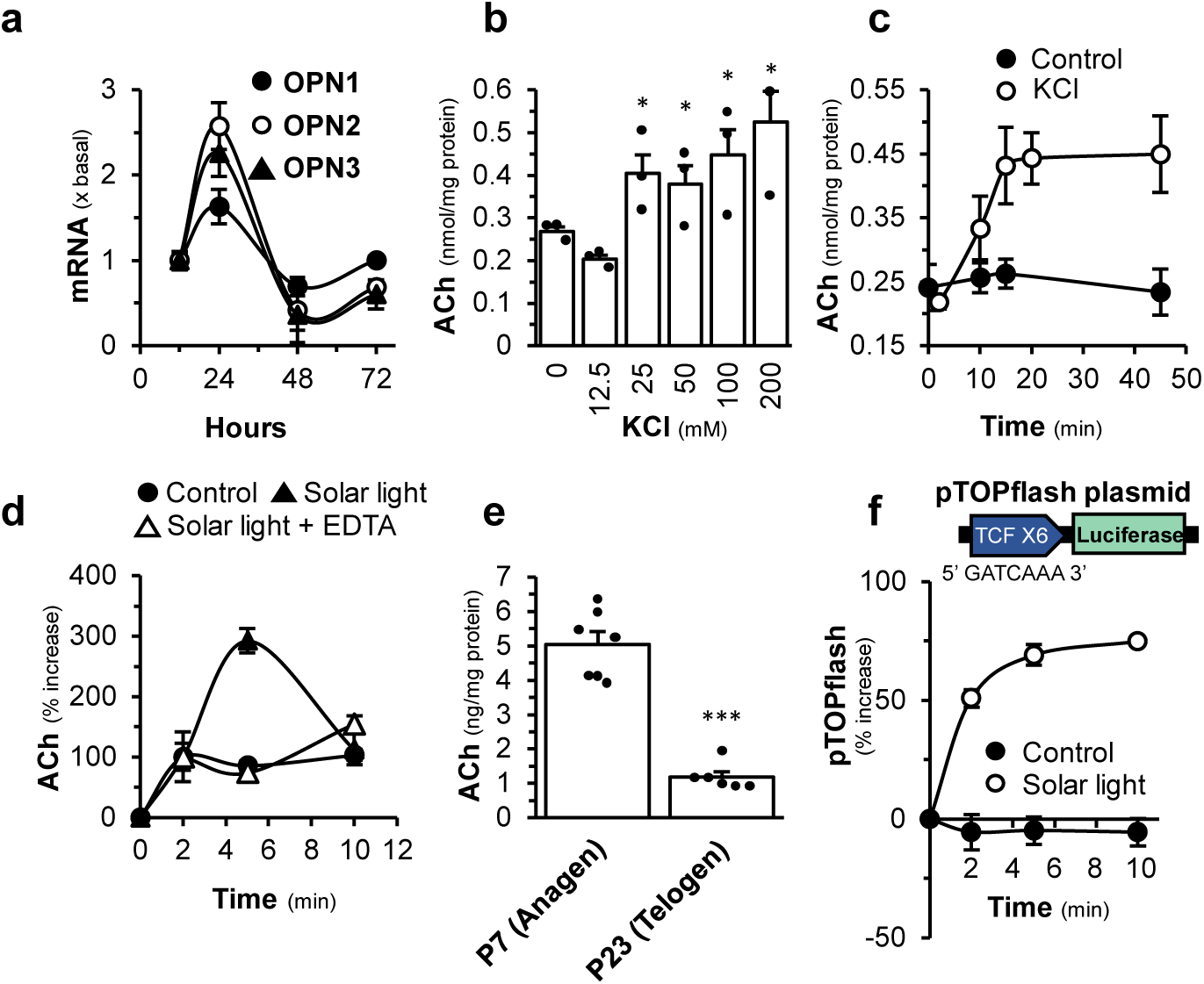
Solar light induces the release of ACh from DPC. **(a)** The levels of mRNAs encoding OPN1, OPN2, OPN3 of cultured DPC harvested on different days (*n*=3). **(b)** ACh release from DPC with various concentrations of KCl, measured by ELISA at 5 min (*n*=3). **(c)** ACh release from DPC over time with 100 mM KCl, measured by ELISA (*n*=3). **(d)** ACh release from DPC over time with the treatment of solar light with or without EDTA (10 mM), measured by LC-MS/MS analysis (*n*=4). **(e)** ACh levels from skin of wild-type mice in anagen (P7) or telogen (P23), measured by LC-MS/MS analysis, normalized by the protein concentration (*n*=6). **(f)** The luciferase activity assay of pTOPflash-transfected DPC was performed after 6 hours of exposure to solar light for various durations (*n*=4). Data are normalized and expressed as the fold of basal value, in comparison to control, in mean ± SEM, * *p* < 0.05, ** *p* < 0.01.

### Cholinergic signalling in hair growth

The immunostaining analysis of cryosections of mouse skin showed the localization of AChE and BChE in dermal papilla, indicated by SOX2 staining, at anagen phase of hair growth cycle, which however disappeared in telogen phase (**Fig. 3a&b**). Thus, AChE and BChE might function robustly during the anagen phase. Because of high expressions of the enzymes, the dermal papilla could be the centre of cholinergic signalling in hair follicle. As expected, AChE and BChE were recognized and localized on the DPC membrane (**Fig. 3c**). To support the role of cholinergic signalling in hair growth, the downstream targets of Wnt signalling, i.e., axin-related protein, (AXIN2), β-catenin, insulin-like growth factor-1 (IGF-1) and ALP, were measured in the present of BW284c51, an AChE inhibitor, for 48 hours in cultured DPCs: the mRNA levels of AXIN-2, IGF-1 and ALP were increased significantly in dose-dependent manners (**Fig. 4a**). Moreover, the growth biomarker of DPC, the mRNA encoding ALP, was increased significantly by a maximum of ∼2.5 folds. The mRNA level of IGF-1, a hair growth promoting factor, was increased by over 3 folds upon BW284c51 treatment. The recombinant Wnt3a served as a positive control here, as an agonist in activating the downstream genes of Wnt (**Fig. 4a**). On the other hand, the role of Wnt/β-catenin signalling in regulating the cholinergic molecules was determined. Valproic acid (VPA), an inhibitor of GSK-3β, activates Wnt/β-catenin signalling in cultured DPC. The mRNA expression of AChE was decreased by ∼80% under the treatment of VPA, and BChE mRNA was increased by ∼3 folds (**Fig. 4b**). The mRNA expression of ChAT was relatively unchanged. Bt_2_-cAMP was served as a control in regulating AChE, BChE and ChAT expressions.

**Figure 3.**
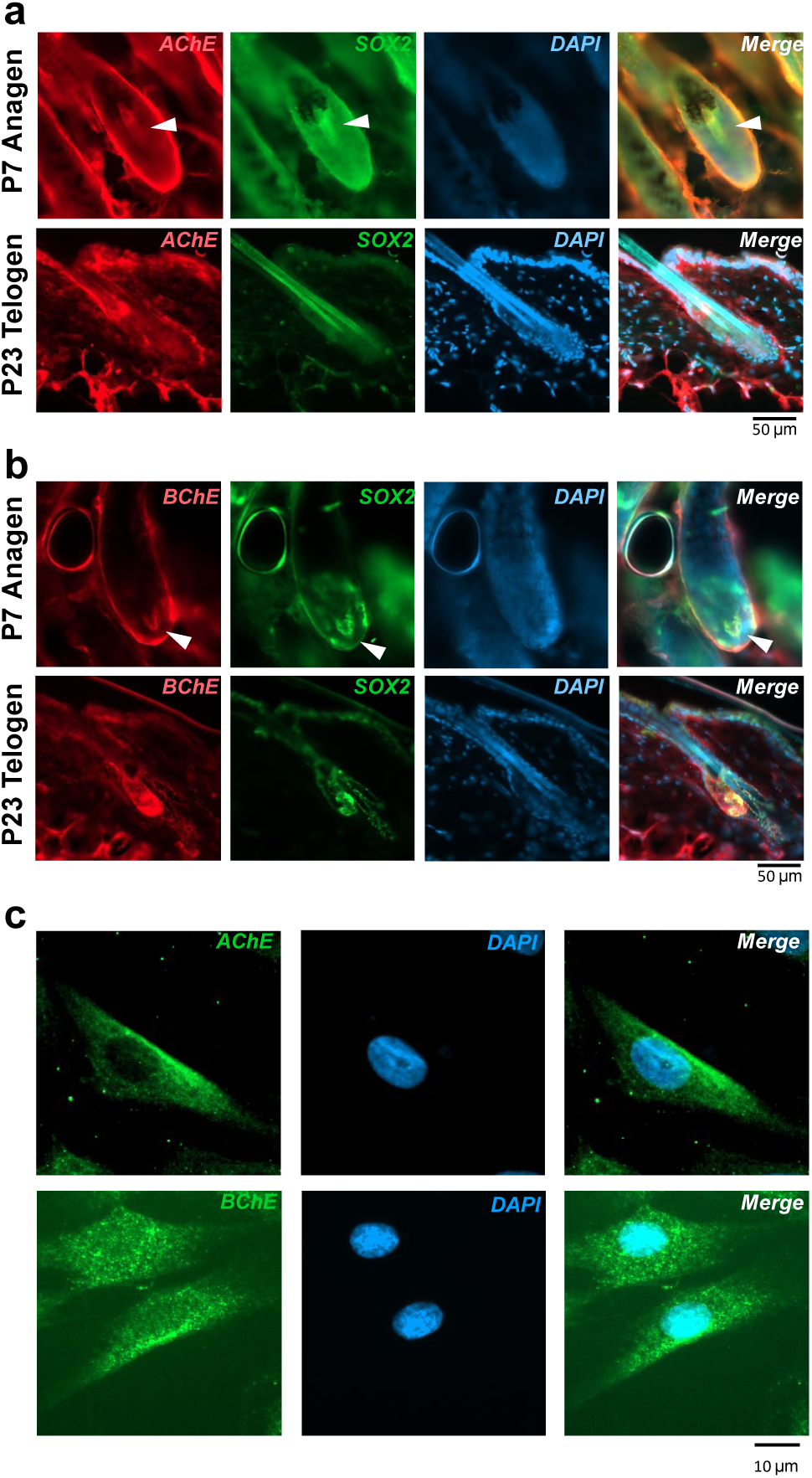
The immunostaining of AChE and BChE in skin from mice and DPC culture**. (a)** Skin from wild-type mice in anagen (P7), telogen (P23) was harvested, fixed, and stained with anti-AChE (*n*=4). **(b)** Skin from wild-type mice in anagen (P7), telogen (P23) was harvested, fixed, and stained with anti-BChE (*n*=4). **(c)** Immunocytochemical staining of AChE (green), BChE (green) and DAPI staining (blue) in DPC, (*n*=4).

**Figure 4.**
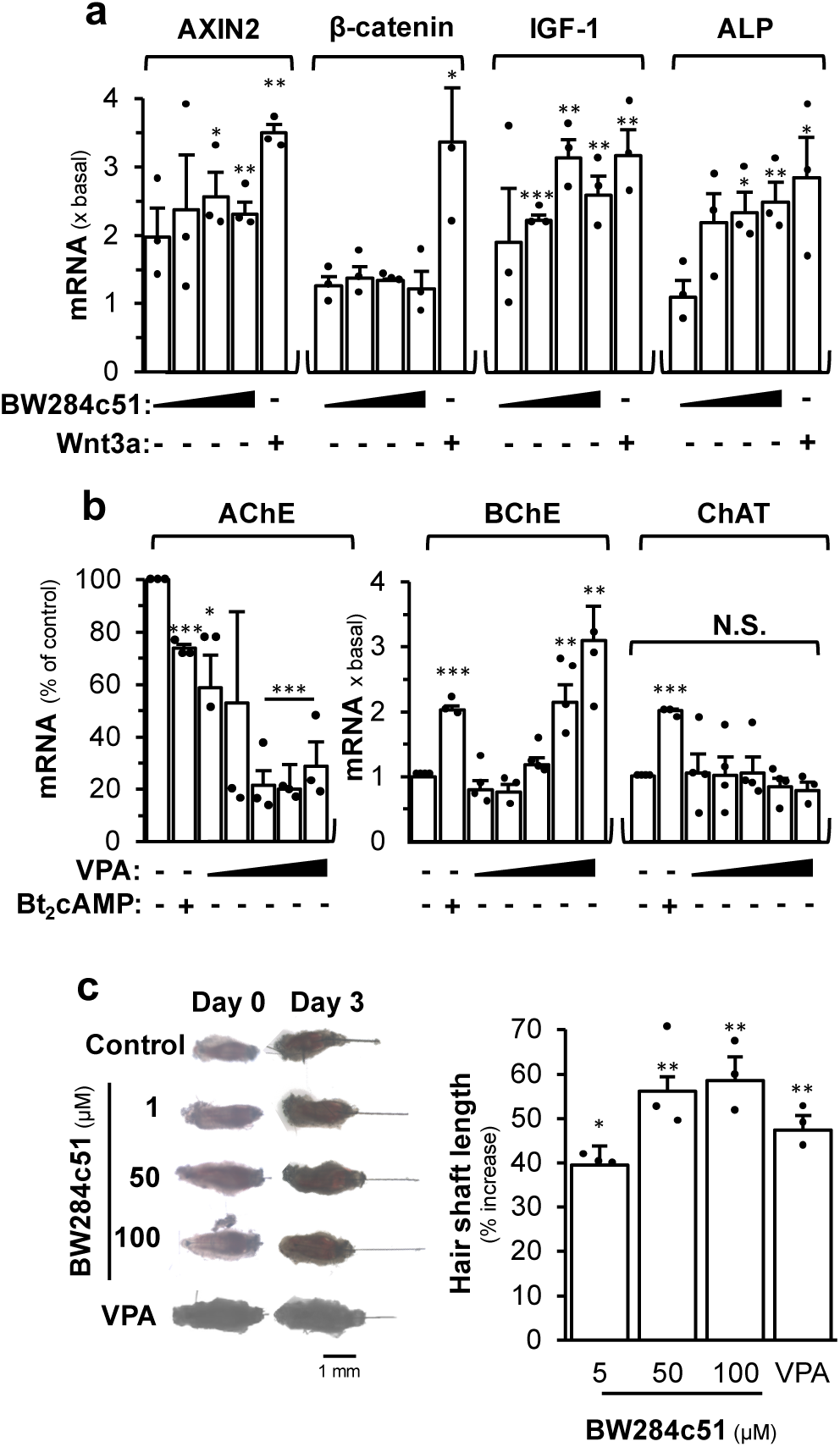
Inhibition of AChE promotes hair growth. **(a)** Cultured DPC was treated with different doses of BW284c51 (25, 50, 100 and 200 µM, as indicated), or Wnt3a (200 ng/mL) 48 hours. The mRNA levels of AXIN2, LEF-1, β-catenin, IGF-1, and ALP are shown (*n*=3). **(b)** Cultured DPC was treated with different doses of VPA (312, 625, 1250, 2500 and 5,000 µM, as indicated), or Bt_2_cAMP (1 mM). The mRNA levels of AChE, BChE, ChAT are shown (*n*=4). **(c)** Individual anagen vibrissae were isolated from the upper lip pad of 4-week-old C57BL/6 male mice. The gross image of cultures treated with different doses of BW284c51, or VPA (5 mM) were taken, as indicated for 3 days (left). The measurement of hair shaft elongation was performed (right) (*n*=3). Data are normalized and expressed as the % increase, or fold of basal value, in comparison to control, in mean ± SEM, * *p* < 0.05, ** *p* < 0.01, *** *p* < 0.001.

To determine the role of AChE inhibitor in hair growth, mouse vibrissae *ex vivo* cultures were used. The isolated vibrissae were treated with different concentrations of BW284c51 for 72 hours. As compared to the control, the length of hair, generated from the follicle being treated with BW284c51, was increased significantly in a dose-dependent manner (**Fig. 4c**). The isolated vibrissae treated with 100 mM BW284c51 showed an increase of hair length by ∼60%, as compared to the control. VPA served as a positive control showing an induction of hair length by ∼45%, less than that of the AChE inhibitor (**Fig. 4c**).

### The cholinergic drugs in hair growth

M4 muscarinic acetylcholine receptor (M4 mAChR) was expressed in hair follicle and DPC, as recognized by immunohistochemical staining of its antibody (**Fig. 5a**). M4 mAChR was co-localized with SOX2, a marker of dermal papilla, indicated that M4 mAChR was expressed predominantly at the dermal papilla. The mRNA expressions of M4 mAChR and alkaline phosphatase (ALP) were decreased in the first 48 hours of culture and increased thereafter. The mRNA expression of choline acetyltransferase (ChAT) remained relatively unchanged during the growth period (**Fig. 5b**). Bethanechol is a non-specific mAChR agonist. Here, bethanechol was applied in pTOPflash-transfected DPC, the luciferase activity was markedly induced in a dose-dependent manner after the treatment: the maximal induction was at ∼5 mM bethanechol (**Fig. 5c**). In addition, bethanechol was applied in cultured DCPs. The mRNA levels of the Wnt-induced signalling downstream genes, i.e., AXIN2, LEF-1, β-catenin, IGF-1 and ALP, were revealed by RT-PCR after the treatment. The results indicated that the levels of mRNAs encoding AXIN2, LEF-1 and β-catenin were significantly increased in a dose-dependent manner upon treatment with bethanechol (**Fig. 5d**). In parallel, the growth markers of DPC, e.g., IGF-1 and ALP, were increased.

**Figure 5.**
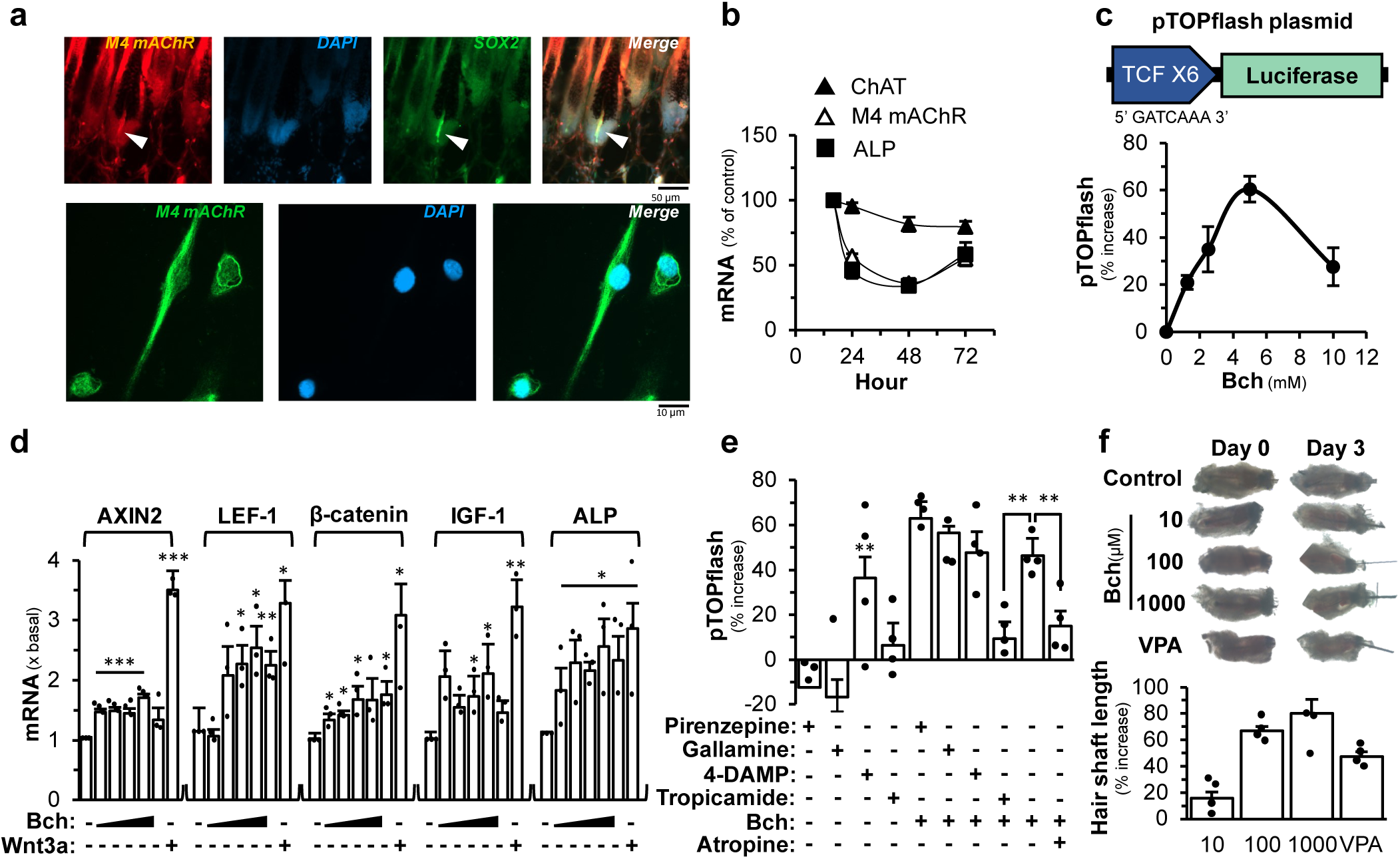
Activation of M4 mAChR promotes hair growth through Wnt signalling. **(a)** Immunohistochemical staining of M4 mAChR (red), DAPI staining (blue), SOX2 (green) in the skin section from wild-type mice in anagen (P7) (upper). Immunocytochemical staining of M4 mAChR (red) and DAPI staining (blue) in DPC (lower). Arrowhead indicates dermal papilla (*n*=3). **(b)** The mRNA levels of ChAT, M4 mAChR and ALP during the DPC growth (*n*=4). **(c)** The luciferase activity assay of pTOPflash-transfected DPC treated with different doses of bethanechol for 24 hours (*n*=4). **(d)** Cultured DPC was treated with different doses of bethanechol (312, 625, 1250, 2500 and 5,000 µM, as indicated), or Wnt3a (200 ng/mL), for 48 hours. The mRNA levels of AXIN2, LEF-1, β-catenin, IGF-1, and ALP are shown (*n*=3). **(e)** The luciferase activity assay of pTOPflash-transfected DPC pre-treated, or treated alone, with pirenzepine (200 µM), or gallamine (200 µM), or 4-DAMP (200 µM), or tropicamide (200 µM), or atropine (50 µM) for 30 min before bethanechol (5 mM) for 24 hours (*n*=4). **(f)** The gross image of individual anagen vibrissae treated with different doses of bethanechol, or VPA (5 mM) for 3 days were taken (upper). The measurement of hair shaft elongation was performed (lower) (*n*=4). Data are normalized and expressed as the % increase, % of control, or fold of basal value, in comparison to control, in mean ± SEM, * *p* < 0.05, ** *p* < 0.01, *** *p* < 0.001.

Pirenzepine (a M1 selective antagonist), gallamine (a M2 selective antagonist), 4-DAMP (a M3 selective antagonist), tropicamide (a M4 selective antagonist), and atropine (an antagonist of muscarinic receptors) were used here to identify the receptor specificity in responding to challenge of bethanechol. In pTOPflash-transfected DPC, the treatment with pirenzepine, gallamine, tropicamide did not induce luciferase activity. In DPC-treated with bethanechol, the luciferase activity was increased by ∼50% (**Fig. 5e**). The pre-treatments with tropicamide and atropine were able to inhibit markedly the bethanechol-induced luciferase activity, suggesting the involvement of M4 mAChR. In line with the notion, the treatment of bethanechol in cultured hair follicles was able to induce hair growth in a dose-dependent manner: the maximal induction by over 80% was at 1 mM bethanechol (**Fig. 5f**). VPA served as a positive control.

GSK3β plays a crucial role in regulating Wnt signalling. GSK3 phosphorylates β-catenin, leading to its destabilization and subsequent degradation, thereby maintaining a low level of β- catenin in the cytosol and nucleus. Treating bethanechol in DPC was able to induce GSK3β phosphorylation at 5 min of incubation, while the pre-treatment of atropine or tropicamide was able to block the induction mediated by bethanechol (**Fig. 6a**). Besides, the GSK3β phosphorylation, induced by bethanechol, was in a transient manner, having a maximal phosphorylation at ∼5 min (**Fig. 6b**). In contrast, VPA induced the phosphorylation at a rather substantial manner. In order to further illustrate the role of M4 mAChR in Wnt signalling, the siRNA knock out of the receptor was performed. The incorporation of siRNA of M4 mAChR in DPCs was able to reduce the receptor expression to ∼60% of the control transfected with negative control siRNA (**Fig. 6c**). In parallel, the protein expression of M4 mAChR in the siRNA transfection was reduced by ∼80% (**Supplementary Fig. 1e**). In the siRNA-transfected cultures, the levels of mRNAs encoding AXIN2, LEF-1, β-catenin and IGF-1, as induced by applied bethanechol, were significantly decreased in a dose-dependent manner upon an increasing dose of siRNA-M4 mAChR (**Fig. 6c**). The results suggested M4 mAChR could be a regulator of Wnt downstream signalling in DPC for hair growth.

**Figure 6.**
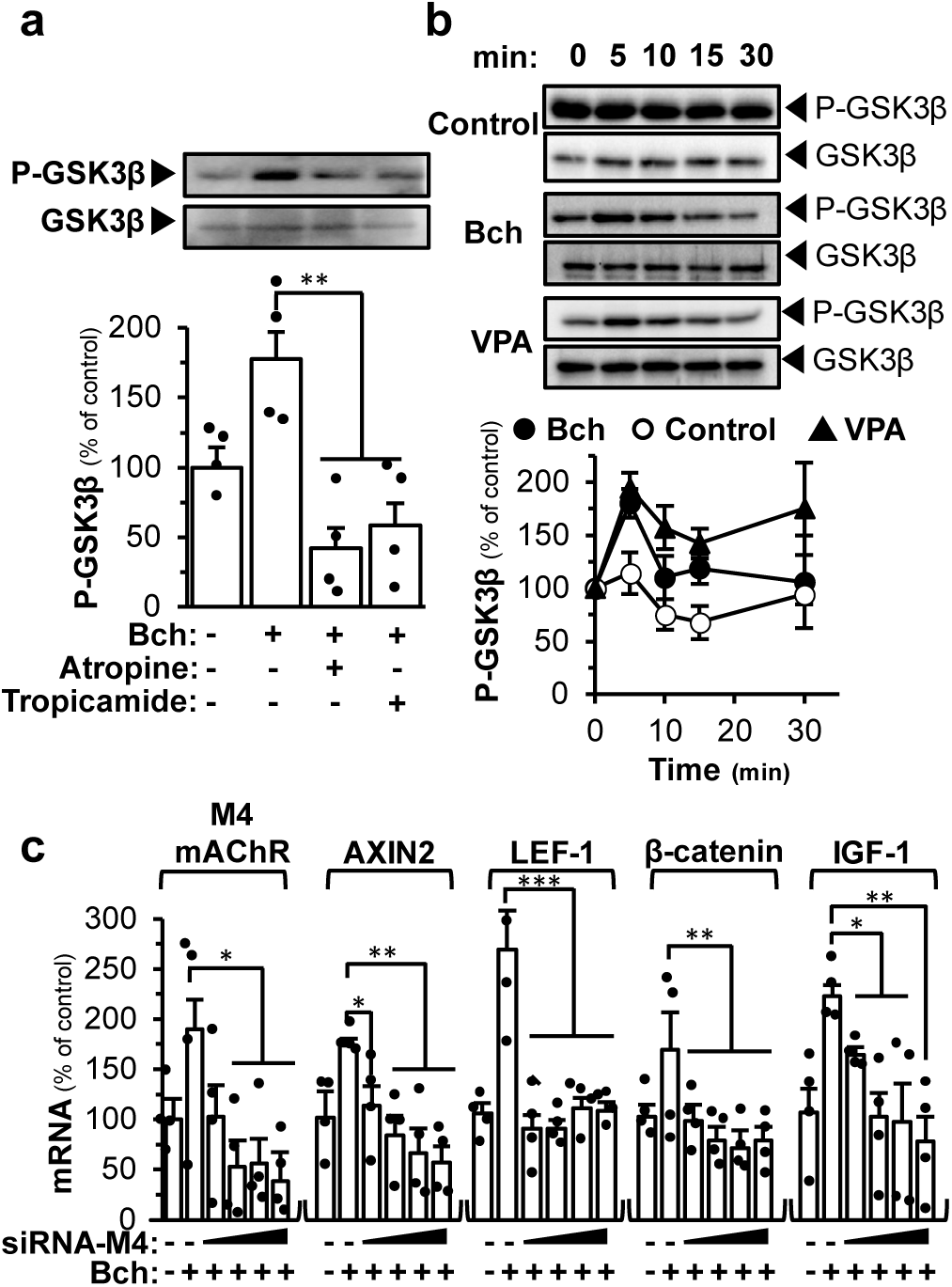
Mechanism of M4 mAChR in activating Wnt/β-catenin signalling. **(a)** Western blot analyses of phosphorylated GSK3β (P-GSK3β) and total GSK3β from DPC treated with bethanechol (5 mM) for 5 min, or preincubation with atropine (50 µM) or tropicamide (200 µM) for 30 min (*n*=4). **(b)** Western blot analyses of phosphorylated GSK3β (P-GSK3β) and total GSK3β from DPC that were subjected to bethanechol (5 mM) or VPA (5 mM) incubation for different times as indicated (*n*=4). **(c)** Different doses of siRNA-M4 (1.25, 2.5, 5, 10 nM, as indicated) were transfected in DPC. The siRNA-M4 transfected DPC were treated with or without bethanechol (5 mM) for 24 hours. The mRNA levels of AXIN2, LEF-1, β-catenin, IGF- 1, and ALP are shown (*n*=4). Data are normalized and expressed as the % of control in comparison to control, in mean ± SEM, * *p* < 0.05, ** *p* < 0.01, *** *p* < 0.001.

To test the cholinergic system being involved in hair growth, the dorsal back of shaved mice was applied every day with the vehicle (negative control), 0.2 mM BW284c51, 25 mM bethanechol and 500 mM VPA (a positive control). Here, the treatments of BW284c51, bethanechol and VPA exhibited an accelerated re-entry of anagen, as determined by hair growth in surface view and H&E cross-section (**Fig. 7a&b**). By histologic analysis, the dermis thickness and the hair weight, and the number of hair follicles were significantly increased under the treatments of BW284c51, bethanechol and VPA (**Fig. 7c-e**). By immunostaining analysis, the expression level of Ki67, a marker of cell proliferation in DPC, was significantly increased, at least by over 150%, in the hair matrix of follicles treated with BW284c51, bethanechol or VPA (**Fig. 7f&g**).

**Figure 7.**
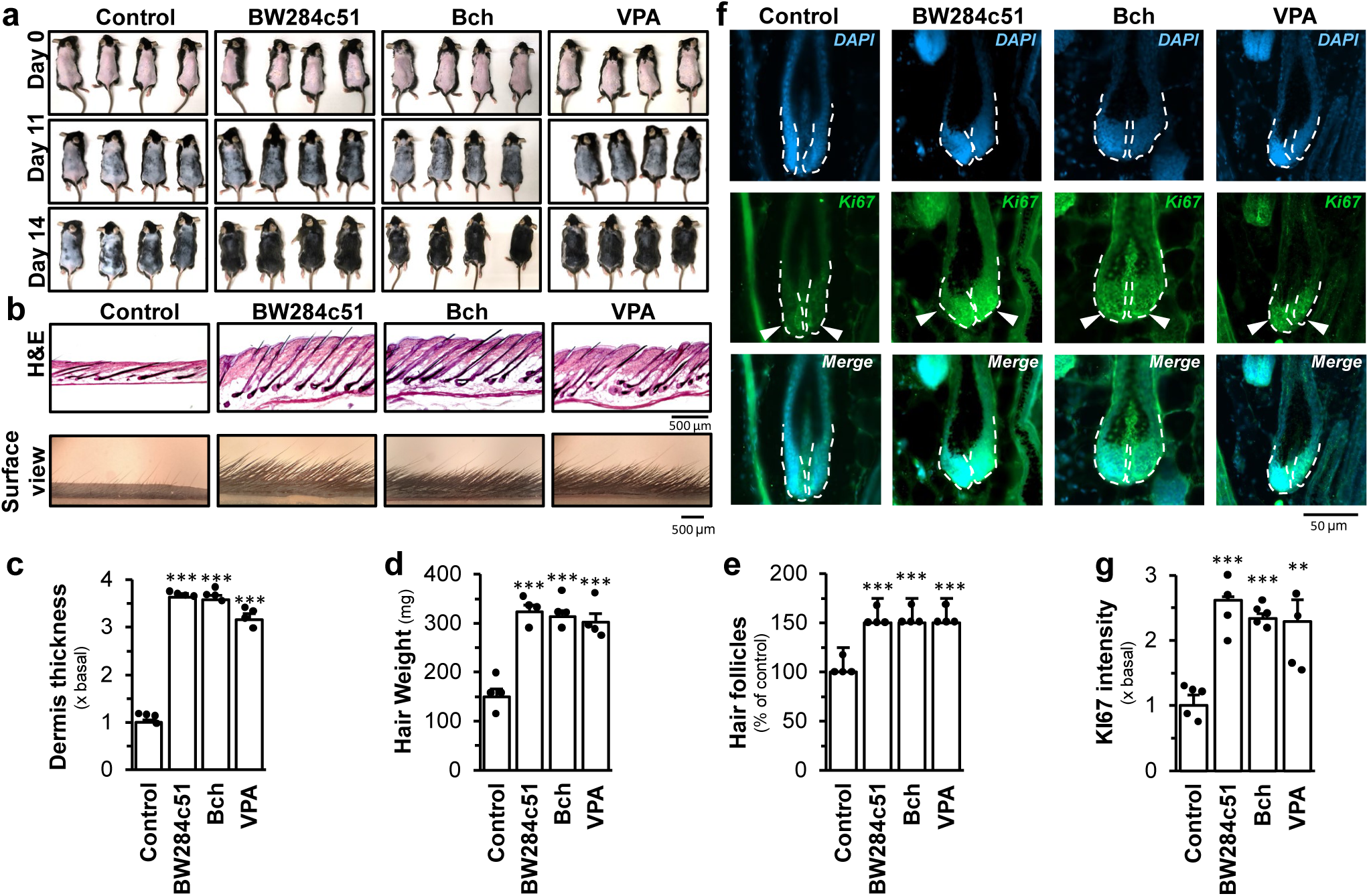
Topical effect of cholinergic drugs accelerated the re-entry of anagen in wild-type mice. Seven weeks old wild-type mice were shaved and treated daily with either topical application of vehicle control, BW284c51 (200 µM), bethanechol (25 mM), VPA (500 mM). **(a)** Representative gross image of the dorsal back of the mice taken at day 0, 11 and 14 (*n*=4). **(b)** Skin was harvested after 14 days and stained with hematoxylin and eosin (H&E) (upper). Surface view shows the status of hair shaft of different treatment (lower) (*n*=4). Measurements of **(c)** dermis thickness (*n*=4), **(d)** hair weight (*n*=4), **(e)** hair follicle number of mouse skin corresponding to different treatments from (a) (*n*=4). **(f)** Immuno-histochemical staining of Ki67 in the hair bulbs of the skin from mice corresponding to different treatments in (a) (*n*=4). **(g)** Quantitative analyses of immunohistochemical staining of Ki67 shown in (f) (*n*=4). Data are normalized and expressed as the % of control, or fold of basal value, in comparison to control, in mean ± SEM, * *p* < 0.05, ** *p* < 0.01, *** *p* < 0.001.

### Down**s**t**r**eam signalling of M4 mAChR in hair growth

The M4 mAChR couples to G_i/o_ protein leading to the inhibition of cAMP production by inhibiting adenylyl cyclase(Van der Westhuizen et al, 2021). The treatment of forskolin activates adenylyl cyclase to increase the endogenous cAMP level (Wang et al, 2022). By pre-treating pGloSensor-22F^TM^ plasmid (a live-cell assay for cAMP signalling coupling to G protein), or pCRE-Luc (a construct for cAMP activation), transfected DPC with forskolin, the increased luminescence, or luciferase signal, was inhibited by bethanechol in a dose-dependent manner (**Fig. 8a&c**). The pre-treatment with tropicamide was able to block the inhibition mediated by bethanechol, which further confirmed the involvement of M4 mAChR in DPC (**Fig. 8b&d**).

**Figure 8.**
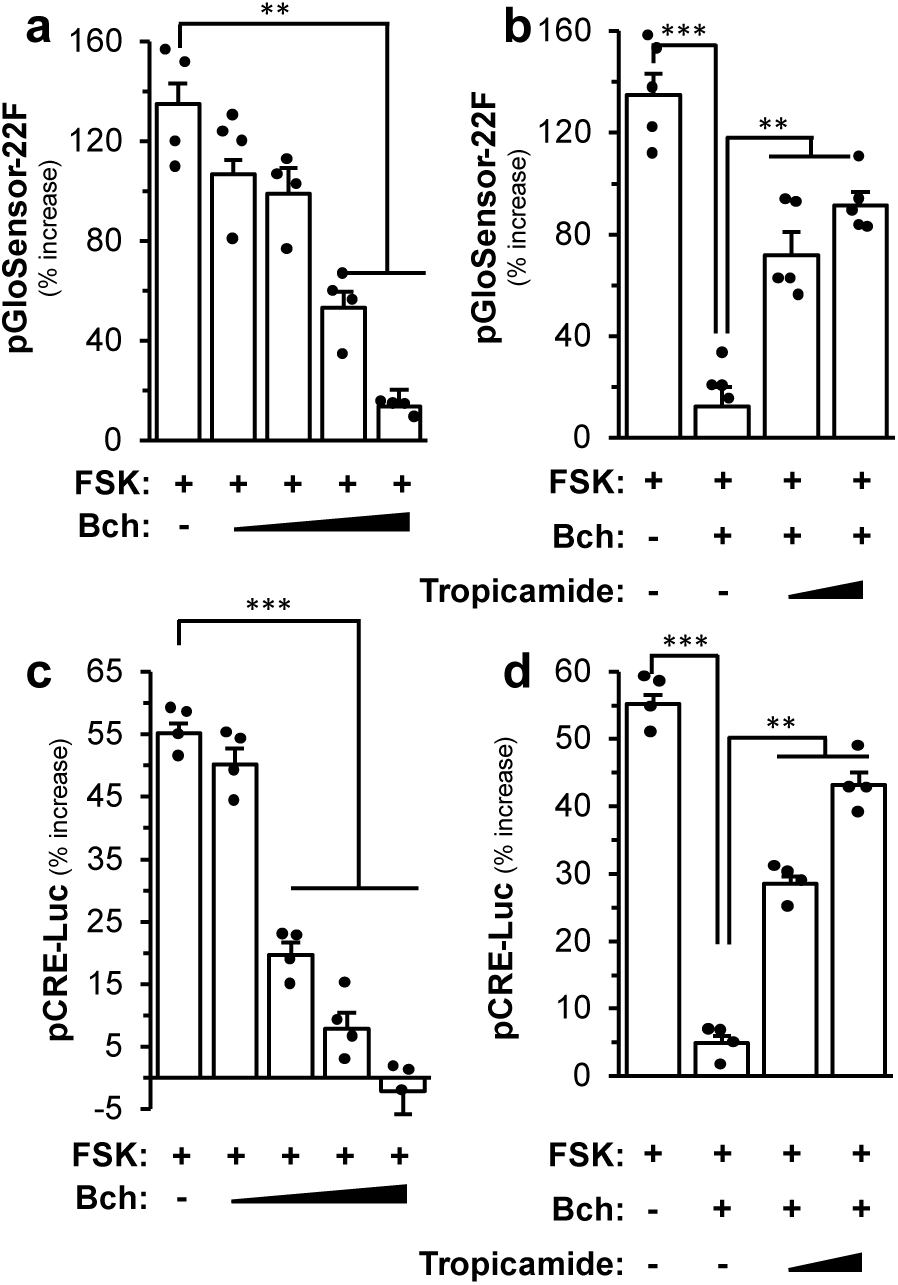
Bethanechol decreases cAMP level in DPC. **(a)** The luciferase activity assay of pGloSensor-22F-transfected DPC treated with different concentration of bethanechol (Bch; 1.25, 2.5, 5, 10 mM, as indicated) in the presence of forskolin (FSK; 10 µM) (*n*=4). **(b)** The luciferase activity assay of pGloSensor-22F-transfected DPC treated with bethanechol (10 mM) in the presence of forskolin (10 µM), either with or without tropicamide (250 or 500 µM) (*n*=4). **(c)** The luciferase activity assay of pCRE-Luc-transfected DPC treated with different concentration of bethanechol (1.25, 2.5, 5, 10 mM, as indicated) in the presence of forskolin (10 µM) (*n*=4). **(d)** The luciferase activity assay of pCRE-Luc-transfected DPC treated with bethanechol (10 mM) in the presence of forskolin (10 µM), either with or without tropicamide (250 or 500 µM) (*n*=4). Data are normalized and expressed as the % increase in comparison to control, in mean ± SEM, ** *p* < 0.01.

The interaction between M4 mAChR and Wnt signalling was poorly studied. We hypothesized that M4 mAChR alone can bypass the Frizzled and LRP5/6 receptors to activate the Wnt signalling in DPC. In pTOPflash-transfected DPC, the application of Wnt3a, or bethanechol, induced the promoter activity (**Fig. 9a**). The activation triggered by Wnt3a was fully blocked by DKK-1 (an inhibitor of Wnt signalling binding to the LRP6 co-receptor) but not by tropicamide. As expected, tropicamide application blocked the activation by bethanechol but not by DKK-1. This implied that there was no interference from the two antagonists. The co-treatment of Wnt3a and bethanechol increased the luciferase activity by ∼100%, higher than the single drug challenge. With the pre-treatment of DKK-1, or tropicamide, in pTOPflash-transfected DPC, the luciferase activity was inhibited partially in the co-treatment of Wnt3a together with bethanechol (**Fig. 9a**). Similar results were further illustrated in pTOPflash-transfected PC12 cells, a neuronal cell line (**Supplementary Fig. 1f**). The co-treatment of Wnt3a and bethanechol induced the phosphorylation of GSK-3β by ∼3 fold, higher than using the single drug. The pre-treatment of DKK-1, or tropicamide, inhibited partially the phosphorylation of GSK-3β triggered by the co-treatment (**Fig. 9b**), suggesting the two signalling events should be independent. This result was further illustrated in *ex vivo* cultures of mouse vibrissae. Both Wnt3a and bethanechol stimulated hair growth. The co-treatment of Wnt3a and bethanechol increased the hair shaft length by ∼80%. As expected, the pre-treatment of DKK-1, or tropicamide, partially inhibited the growth triggered by the co-treatment (**Fig. 9c**).

**Figure 9.**
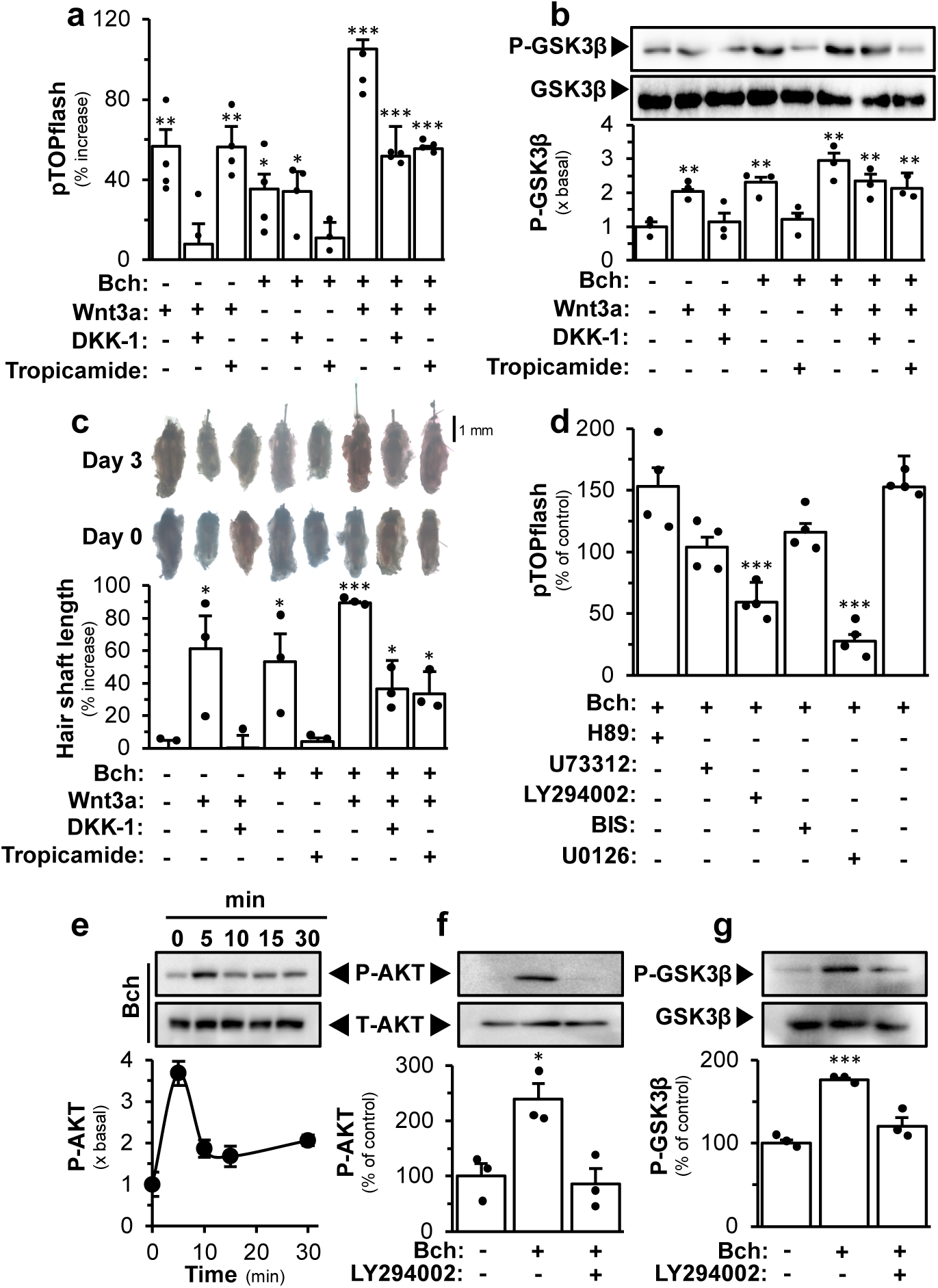
Relationship of signaling cascades of M4 mAChR and Wnt/β-catenin. **(a)** The luciferase activity assay of pTOPflash-transfected DPC pre-treated with or without DKK-1 (300 ng/mL), or tropicamide (500 µM), for 30 min before the treatment of bethanechol (Bch; 5 mM), or Wnt 3a (200 ng/mL), or simultaneously with both treatments, for 24 hours (*n*=4). **(b)** Western blot of phosphorylated GSK3β (P-GSK3β) and total GSK3β from DPC which were subjected to the same treatment in (a) for 5 min (*n*=3). **(c)** The gross image of individual vibrissae subjected to the same treatment in (a) for 3 days were taken (upper). The measurement of hair shaft elongation was performed (lower) (*n*=3). **(d)** The luciferase activity assay of pTOPflash-transfected DPC pre-treated or treated alone with H89 (10 µM), or U73312 (100 nM), or LY294002 (50 µM), or BIS (50 nM), or U0126 (10 µM) for 30 min before the treatment of bethanechol (5 mM) for 24 hours (*n*=4). **(e)** Western blot of phosphorylated AKT (∼60 kDa) and total AKT (∼60 kDa) from DPC under bethanechol (5 mM) incubation for different times (*n*=3). **(f)** Western blot of phosphorylated AKT and total AKT as in (e) for 5 min, or preincubation with LY294002 (50 µM) for 30 min (*n*=3). **(g)** Western blot of phosphorylated GSK3β (P-GSK3β) and total GSK3β from DPC subjected to the same treatment in (f) (*n*=3). Data are normalized and expressed as the % increase, % of control, or fold of basal value, in comparison to control, in mean ± SEM, * *p* < 0.05, ** *p* < 0.01.

**Figure 10.**
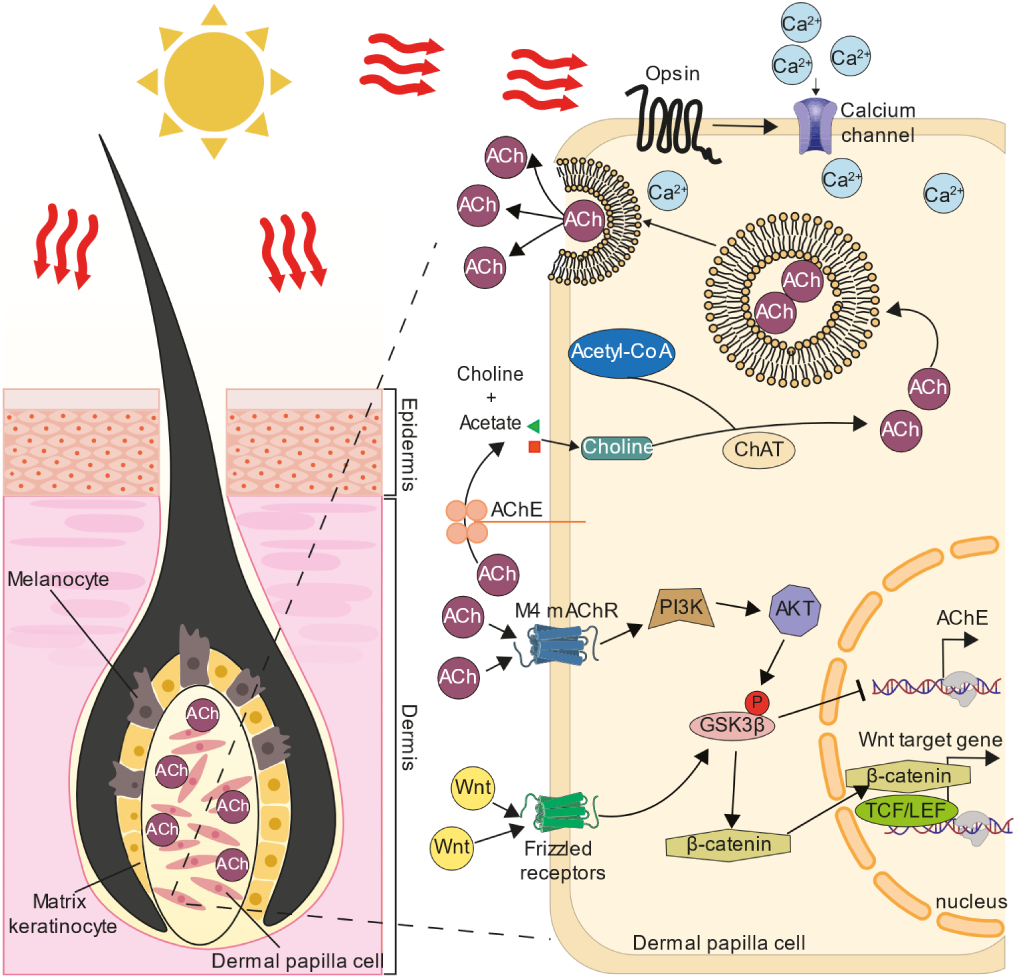
Proposed hypothesis for AChR in hair growth. Under the stimulation of solar light, the influx of Ca^2+^ causes the release of ACh from DPC of hair follicle, and subsequently which acts as an autocrine agent in activating M4 mAChR. The activation of M4 mAChR can bypass the Frizzled and LRP5/6 receptors to activate the Wnt signalling by the phosphorylation of GSK-3β through the PI3K/AKT pathway. Together with the action of secreted Wnt proteins from hair follicle epithelium, the activation of Wnt/β-catenin signalling can be further augmented. The phosphorylated GSK-3β induced CREB DNA binding activity inhibit the expression of AChE, as a positive feedback loop.

To investigate the relationship between mAChR and Wnt signalling pathways, different inhibitors, including H89 (PKA inhibitor), U73312 (PLCβ inhibitor), LY294002 (PI3K inhibitor), BIS (PKC inhibitor), U0126 (ERK/MEK inhibitor), were pre-treated to the cultured DPC, as to block the activation triggered by bethanechol. Treatments of LY294002 and U0126 were able to inhibit the effect of bethanechol, significantly (**Fig. 9d**). U73312 and BIS showed minor inhibition, but not at a significant level. The pathway of PI3K/AKT/mTOR are known to have a direct relationship with Wnt signalling via phosphorylation of GSK3β. Here, the treatment of bethanechol in DPCs induced the phosphorylation of AKT in a time-dependent manner and peaked at 5 min: this effect was fully inhibited by pre-treatment with LY294002 (**Fig. 8e&f**). Moreover, the bethanechol-induced GSK-3β phosphorylation was inhibited by the pre-treatment of LY294002 (**Fig. 9g**). This further verified the AKT could be the mediator between the signalling of M4 mAChR and Wnt pathway.

## DISCUSSION

For the first time, we have given a detailed mechanistic analysis of cholinergic system in regulating hair growth. Under the stimulation of solar light, ACh is being released from DPC, and which thereafter activates M4 mAChR, as well as Wnt signalling pathway, at the DPC (**Fig. 9**). This stimulatory effect of Wnt signalling by M4 mAChR in promoting hair growth is mediated via PI3K/AKT and ERK/MEK signalling pathways. Several lines of evidence support this notion: (i) the expression and localization of cholinergic molecules in DPC; (ii) induction of ACh release by solar light; (iii) activation of M4 mAChR triggering Wnt signalling and hair growth; (iv) blockage of hair growth by mAChR antagonist; and (v) regulation of cholinergic enzyme by Wnt agonist. Our findings support the relationship of hair pigmentation and hypertrichosis in the known therapy of having intake of cholinesterase inhibitors. For an example in Alzheimer’s patients, 24 out of 62 patients reported hair darkening in the occipital region after using cholinesterase inhibitor for at least 6 months (Chan et al, 2020). In another example, a Caucasian 80-year-old male who received AChE/BChE inhibitor rivastigmine for one month was diagnosed acquired localized hypertrichosis on both forearms (Imbernón-Moya et al, 2016). These findings highlight the potential regulation of hair growth by cholinergic system. Having this hypothesis, the cholinergic molecules in hair follicles may be a potential therapeutic target for alopecia. Besides, AChE inhibitors are currently approved by the US Food and Drug Administration for the treatment of Alzheimer’s disease, and in general these drugs are safe to use (McGleenon et al, 1999; Vecchio et al, 2021). To fully comprehend the potential of these inhibitors as an alopecia treatment, additional preclinical and clinical studies are required. Our studies raise the possibility that the cholinergic drug can be a new niche to cure alopecia.

Wnt signalling plays an important role in growth and development of hair follicles. Here, we have demonstrated the activation of Wnt signal in DPC could be triggered by M4 mAChR. The relationship between cholinergic system and Wnt signalling has been reported in various cell types, but the detailed mechanism is not clear (Jensen et al, 2012; Labed et al, 2018; Renz et al, 2018). Huperzine A, a selective AChE inhibitor, has been shown to inhibit GSK3α/β activity and enhance the level of β-catenin in transgenic mouse model of Alzheimer’s disease, as well as in cultured SH-SY5Y cells (Wang et al, 2011). In parallel, the application of Donepezil in scopolamine-induced mice showed similar inhibition on the activity of GSK3α/β (Machhi et al, 2016). Nor-galanthamine, an alkaloid and a potent AChE inhibitor, triggered the anagen-activating signalling pathways, including ERK1/2, AKT and β-catenin pathway, in cultured DPC (Yoon et al, 2019). These findings are consistent with our results that the inhibition of AChE in DPC can promote hair growth via activation of AChR and subsequently the Wnt signalling.

ACh was first identified in skin at 1962 (Scott, 1962). The functionalities of ACh in skin have been proposed, and one of them is playing a role in skin pigmentation (Arredondo et al, 2003; Wui et al, 2020a). Keratinocyte and melanocyte form a so called “skin synapse” in the skin epidermis. Keratinocyte is able to release ACh, triggered by light-activated opsin, and that acts on mAChR of melanocyte, as such to regulate the production and release of melanin (Wu et al, 2018; Wu et al, 2020b). Similar to the situation in skin, the hair DPC could release ACh, possibly triggered by solar light-activated opsins (Bellono et al, 2014; Kelley & Davies, 2016). In contrast, the released ACh is acting as an autocrine to trigger the responses on DPC itself. However, the exact sub-types of opsins in directing this function are not known; but at least the expressions of opsins 1-3 are peaked at first day after DPC culture. Besides, the level of ACh level is shown to be higher in anagen phase than in telogen phase of hair follicle, which is consistent with the hypothesis that the opsin is interacting with cholinergic molecules via solar light to regulate hair growth.

Although various mAChRs, including M1 to M5, were found in the dermal papilla. Here, we propose M4 mAChR should be the primary target to mediate the cholinergic signalling in hair growth. In line to this notion, M3 and M4 mAChR have been reported to involve in keratinocyte migration and hair growth cycling. The M4 mAChR knock-out mice displayed retardation of hair follicle morphogenesis (Hasse et al, 2007), and similarly a decreased keratinocyte migration was observed upon transfection with siRNA of M4 mAChR (Chernyavsky et al, 2004). The reciprocal effect was revealed in keratinocyte transfected with siRNA-M3 mAChR and M3 mAChR KO mice (Chernyavsky et al, 2004). In line to these reports, an inhibition of M3 mAChR using 4-DAMP was able to stimulate the Wnt signalling in cultured DPCs, as shown here. Conversely, the inhibition of M4 mAChR using tropicamide or M4 mAChR targeted siRNA can inhibit the Wnt signalling, even in the present of bethanechol. Moreover, with the pretreatment of the non-specific antagonist atropine in DPC, Wnt signalling was inhibited even with the induction of bethanechol. Thus, the inhibitory effect of M4 mAChR on Wnt signalling outweighed the stimulatory effect of M3 mAChR in the presence of atropine. This suggests that the role of M4 mAChR may take higher priority than M3 mAChR.

## MATERIALS AND METHODS

### Cell cultures and chemicals

Immortalized DPCs were obtained from Applied Biological Materials (Richmond, BC, Canada). SH-SY5Y cells (a human neuroblastoma line), human keratinocytes (HaCaT) and rat pheochromocytoma PC12 cells were purchased from American Type Culture Collection (ATCC, Manassas, VA). DPC, SH-SY5Y and HaCaT cells were cultured in Dulbecco’s modified Eagle medium supplemented with 10% fetal bovine serum (FBS) and 1% (v/v) penicillin/streptomycin (stock as 10,000 U and 10,000 mg/mL) in 5% CO_2_ at 37 °C. PC12 cells were cultured in DMEM supplemented with 6% fetal bovine serum (FBS), 6% horse serum, and 1% penicillin/streptomycin (10,000 U/mL and 10,000 μg/mL). All culture reagents were purchased from Thermo Fisher Scientific (Waltham, MA). The siRNA-M4 was custom synthesized by GenePharma (Shanghai, China). The target sequences for the human *CHRM4* (NM_000741) gene was 5’-CCT CTA CAC CGT GTA CAT C-3’ as reported previously(Chernyavsky et al., 2004). The sequence of negative control siRNA was 5’-UUC UCC GAA CGU GUC ACG UTT-3’.

### DNA transfection and luciferase reporter assay

The plasmid having TCF/LEF-firefly luciferase reporter (pTOPflash) with two repeats, each containing three copies of the TCF-binding site upstream of thymidine kinase minimal promoter, was purchased from Upstate Biotechnology (Lake Placid, NY). The pGloSensor-22F cAMP plasmid was purchased from Promega (Madison, WI). GloSensor™ luciferase contains cAMP binding domain fused to a circularly permuted form of firefly luciferase. The pCRE-Luc contains CRE promoter sequences with a luciferase reporter, purchased from BD Biosciences. The siRNA-M4, 5’-CCT CTA CAC CGT GTA CAT C-3’ (corresponding to amino acids 259–279) and negative control siRNA, 5’-UUC UCC GAA CGU GUC ACG UTT-3’, was custom synthesized by GenePharm (Shanghai, China), as reported previously (Chernyavsky et al., 2004). The reporter constructs were then transfected using human hair follicle dermal papilla cell (HFDPC) Avalanche® Transfection Reagent (EZ Biosystems, College Park, MD) according to the given instruction. In short, DPC was seeded into 6-well plate incubated overnight in 5% CO_2_ at 37°C in a humidified environment without the use of penicillin/streptomycin in the culture medium. After 3 hours incubation with the transfected mixture, the medium was aspirated and replaced by DMEM supplement with 10% FBS and different drug treatment. For the transfection of pGloSensor-22F cAMP plasmid and siRNA- M4, the transfection mixture was incubated overnight. After 24 hours, the cells were collected for the luciferase reporter assay. The luciferase assay was performed using Pierce^TM^ firefly luciferase kit (Thermo Fisher Scientific). The luminescent reaction was quantified in a GloMax® 96 microplate luminometer (Thermo Fisher Scientific) and normalized to the total protein. For the pGloSensor-22F cAMP assay, the pre-incubated medium was replaced by fresh medium containing 500 µM of 3-isobutyl-1-methylxanthine and treated with or without drug for 15 min before the addition of 10 µM forskolin. After 15 min of forskolin treatment, the culture was lysed, followed by the luciferase assay.

### Sucrose density gradient analysis

Different AChE forms were separated by sucrose density gradient analysis according to previous method (Choi et al., 2008; Xia et al., 2022). In brief, a continuous 5-20% sucrose gradient in lysis buffer containing 10 mM HEPES, pH 7.5, 1 mM EDTA, 1 mM EGTA, 0.2% Triton X-100, and 150 mM NaCl was prepared in a 12-mL polyallomer ultracentrifugation tube. Two hundred μL (1 μg/μL) cell lysates mixed with sedimentation markers, including alkaline phosphatase (6.1 S) and β-galactosidase (16 S), were loaded onto the gradients followed by centrifugation as 38,000 rpm in SW41 Ti Rotor (Beckman, Palo Alto, CA) at 4 °C for 16 hours.

Approximately 48 fractions were collected for determination of AChE activity by Ellman assay. In Ellman assay, 0.1 mM tetra-iso-propylpyrophosphoramide (iso-OMPA; Sigma-Aldrich, St Louis, MO), 0.625 mM acetylthiocholine (ATCh; Sigma-Aldrich), and 0.5 mM 5,5-dithiobis-2-nitrobenzoic acid (DTNB) in 80 mM Na_2_HPO_4_, pH 7.4, was added to the sample. The mixture was incubated at room temperature for 30 min, and AChE activities were measured at 405 nm using Multiskan™ FC microplate photometer. The enzyme activity was expressed as absorbance units/min/g of protein.

### Real-time PCR

Total RNA was extracted from cultured DPCs using RNAzol™ Reagent (Sigma-Aldrich), and 3 μg of RNA was reverse transcribed using PrimeScript™ RT reagent kit (Takara, San Jose, CA), according to manufacturer’s instruction. Template cDNAs were subjected to RT-PCR using the following specific primers provided in **Supplementary Fig. 3**. Real-time PCR was performed in LightCycler 480 (Roche Molecular Biochemical, Indianapolis, IN) using KAPA SYBR FAST qPCR kits in accordance with the manufacturer’s instruction. The 2^ΔΔCt^ method was used to calculate the relative expression levels.

### Vibrissae culture

Mouse vibrissae were carefully isolated from the upper lip pad of 4-week-old C57BL/6 male mice and cultured in Williams E medium (Sigma-Aldrich), supplemented with 1% (v/v) penicillin/ streptomycin solution, under 5% CO_2_ at 37 °C. In each experiment, follicles were collected from a single mouse and assigned to individual treatment groups. Each treatment group consisted of four hair follicles. Vibrissae were incubated in various conditioned medium for 3 days, and the increase in length of hair was measured from days 0 to 3.

### Animals and *in vivo* hair growth test

Six-week-old male C57BL/6 mice were chosen for the experiment and allowed to adapt to their new environment for 1 week. The hairs on the backs of 7-week-old mice, whose hair follicles were in the telogen phase, were shaved with a hair clipper. Various drugs were applied topically daily for up to 14 days, with four mice in each group. All reagents used for the hair re-growth test were dissolved in a vehicle composed of 100% ethanol and mixed with equal amount of cream.

### Skin cryosection preparation

The dorsal skin from C57BL/6 mice was shaved to remove hair and collected after being sacrificed. Skin tissue was fixed with 4% paraformaldehyde for 1 hour at room temperature and then perfused with 30% sucrose solution in PBS. Afterward, the skin was embedded in OCT and frozen at-80 °C overnight followed by cutting into 30 μm section with Thermo CryoStar NX 70 Cryostat (Thermo Fisher Scientific). The section was preserved at-20 °C for the following experiments.

### Immuno-fluorescent staining

Cultured cells were grown on glass coverslips in 35-mm culture dishes. After PBS wash, cells were fixed with 4% paraformaldehyde for 15 min. Cells were incubated with or without 0.1% Triton X-100 in PBS for 10 min and then blocked by 5% BSA for 1 hour. Cultures were stained with primary antibodies for 16 hours at 4 °C, followed by Alexa 488/555/647-conjugated secondary antibodies (Sigma-Aldrich) for 2 hours. The following antibodies were used: anti-AChE at 1:100 (Santa Cruz), anti-BChE at 1:100 (R&D System), anti-M4 mAChR at 1:100 (Abcam Ltd), anti-ChAT at 1:100 (Abcam Ltd), anti-SYP at 1:100 (Santa Cruz), anti-SOX2 at 1:100 (Abcam Ltd), anti-Ki67 at 1:100 (Abcam Ltd). Samples were mounted with ProLong™ Gold antifade mountant with or without DAPI (Thermo Fisher Scientific). Samples were then examined by Zeiss Celldiscoverer 7 automated microscope (Zeiss, Oberkochen, Germany).

### Measurement of ACh

The amount of ACh in conditioned medium or skin tissue from wild-type mice was measured using Acetylcholine ELISA Kit (Colorimetric) (Abcam Ltd) and LC–MS/MS. Briefly, cells were seeded and cultured in a 60-mm culture dish. After 48 hours, the culture medium was replaced by DMEM without FBS supplied with 20 µM BW284c51 and 20 µM iso-OMPA for 30 min. The culture medium was replaced by Ringer’s buffer (7.2 g/L NaCl, 0.37 g/L KCl, 0.17 g/L CaCl_2_, and pH 7.4) with 20 µM BW284c51 and 20 µM iso-OMPA, then cells were exposed under the solar light simulator machine (Newport Corporation, Irvine, CA) or treated with KCl. The experimental irradiance contained UVA (0.5 mW/m2), UVB (50 mW/m2), and UVC (50 mW/m^2^). The Ringer’s buffer containing ACh, released by cells, was collected, heated at 55 °C for 5 min to deactivate cholinesterase and concentrated with freeze drying, followed by ACh assay performed. In brief, the amount of choline in each sample was determined using the probe for choline before and after adding AChE. The amount of ACh was acquired by subtracting the amount of choline before adding AChE from that after adding AChE. For the ACh amount in the skin tissue, the mouse skin tissue was heated at 55 °C for 5 min to deactivate cholinesterase immediately after dissection in lysis buffer containing 10 mM HEPES, pH 7.5, 1 mM EDTA, 1 mM EGTA, 0.2% Triton X-100, and 150 mM NaCl. The samples were then homogenized, vortex and centrifuge to obtain the extract. For the LC-MS/MS analysis, LC-MS/MS analysis was conducted an Agilent 6410B triple quadrupole mass spectrometer (QQQ-MS/MS) with ESI source (Agilent Technologies, Waldbronn, Germany) coupled with Agilent 1290 Infinity Binary Liquid Chromatography Systems. Chromatographic separation was achieved on a PC HILIC (150 mm × 2 mm, 3 µm, Osaka Soda, Osaka, Japan) at 35 °C. For conditioned medium, after freeze drying, the solid obtained was redissolved in 100% acetonitrile before injection into the mass spectrometer. For tissue sample, an equal amount of acetonitrile was added to the sample to precipitate the protein before injecting into the mass spectrometer. The mobile phase consisted of water containing 20 mM ammonium formate and 0.2% formic acid (A) and acetonitrile containing 0.1% formic acid (B) at a flow rate of 0.4 ml/min. The gradient elution was programmed as follows: 0–2 min, 2% A; 2-2.01 min, 2-25% A; 2.01–4 min, 25% A; 4–5 min, 25-50% A; 5-8 min, 50% A; 8–10 min; 50-2% A and 10-13 min, 2% A.

### Statistics analysis

Each result is represented as the mean ± SEM, calculated from independent replicates. Comparisons of the means for untreated control cells and treated cells were analysed using Student’s *t*-test. Significant values were represented as \**p* < 0.05, \*\**p* < 0.01, \*\*\**p* < 0.001.

## DATA AVAILABILITY STATEMENT

All data needed to evaluate the conclusions in the paper are present in the paper and/or the Supplementary Materials.

## ACKNOWLEDGMENTS

This work is supported by supported by Hong Kong Research Grants Council Hong Kong (GRF 16100921); Zhongshan Municipal Bureau of Science and Technology (2019AG035); Guangzhou Science and Technology Committee Research Grant (GZSTI16SC02; GZSTI17SC02); GBA Institute of Collaborate Innovation (GICI-022); The Key-Area Research and Development Program of Guangdong Province (2020B1111110006);Special project of Foshan University of Science and Technology in 2019 (FSUST19-SRI10); Hong Kong RGC Theme-based Research Scheme (T13-605/18-W); Hong Kong Innovation Technology Fund (ITS/500/18FP; MHP/004/21; GHP/016/21SZ; ITCPD/17-9; ITC- CNERC14SC01); TUYF19SC02, PD18SC01 and HMRF18SC06; HMRF20SC07, AFD20SC01; Shenzhen Science and Technology Innovation Committee (ZDSYS201707281432317).

## AUTHOR CONTRIBUTIONS

G.K.W.Y., Conceptualization, Methodology, Investigation, Visualization, Writing—original draft, Writing—review & editing; K.W.L., Methodology, Investigation; S.Y.J.X., Methodology; K.Q.Y.W., Investigation; A.X.G, Investigation; Q.W.S.L., Investigation; J.Y.M.H., Software H.C.T.C., Investigation; M.S.S.G., Investigation; K.W.K.T., Supervision, Writing—review & editing.

---

**Supplementary Fig. 1.**
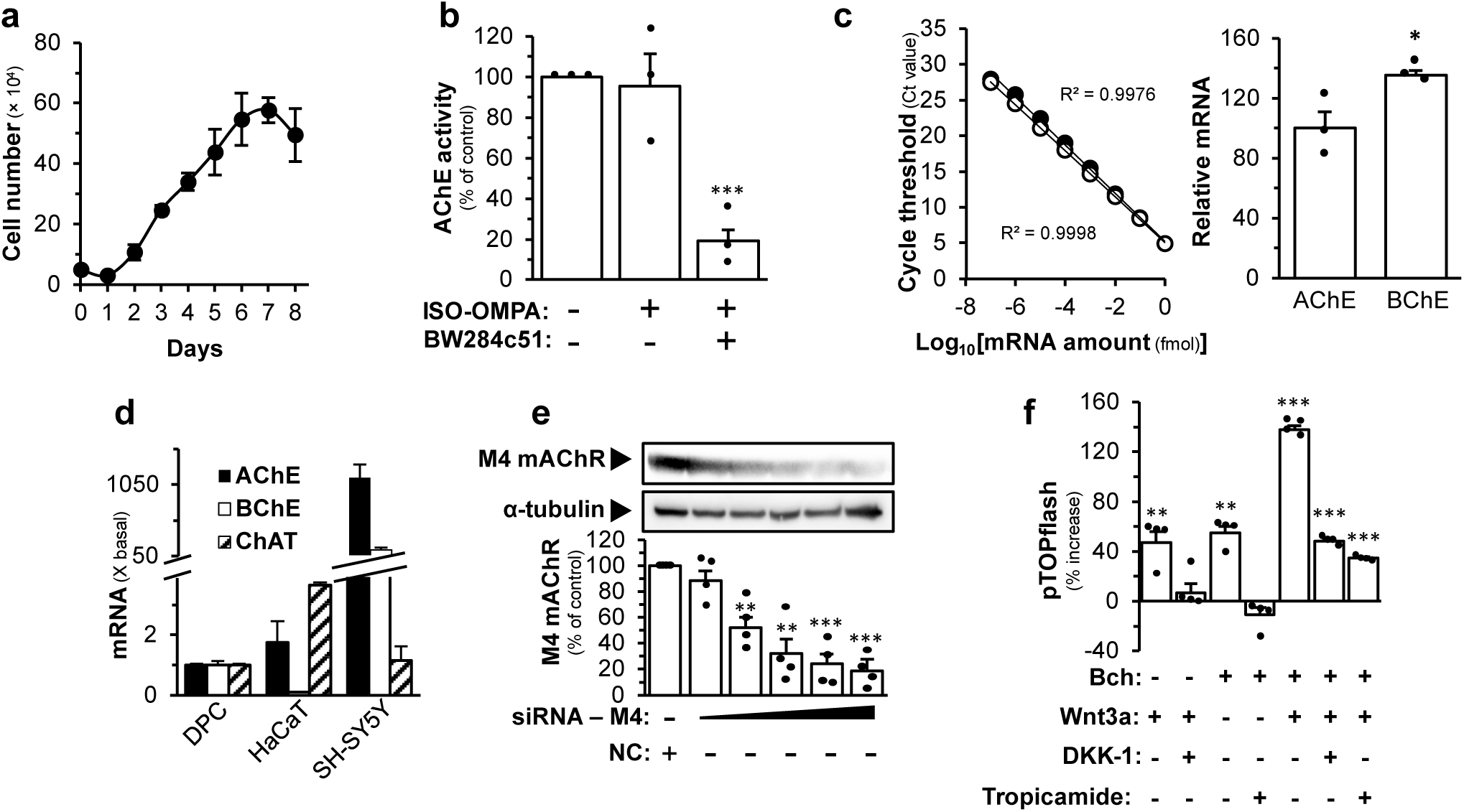
Cholinergic molecules in Wnt signaling. **(a)** Growth curve analysis of DPC (*n*=4). **(b)** AChE activity of DPC treated with or without iso-OMPA (50 µM) or BW284c51 (50 µM) (*n*=3). **(c)** The standard curve of AChE and BChE mRNA levels was obtained by quantitative PCR (left). The mRNA levels of AChE and BChE in DPC cultures were determined by the absolute quantitative PCR (right) (*n*=3). **(d)** The mRNA levels of AChE, BChE and ChAT are shown in cultured DPC, HaCaT and SH-SY5Y (*n*=4). **(e)** Western blot of M4 mAChR (∼55 kDa) and α-tubulin (∼50 kDa) from DPC transiently transfected with siRNA- M_4_ (0.625, 1.25, 2.5, 5, 10 nM, as indicated) or negative control (10 nM) (*n*=4). **(f)** The luciferase activity assay of pTOPflash-transfected PC12 pre-treated with or without DKK-1 (300 ng/mL) or tropicamide (500 µM) for 30 min before the treatment of Bch (5 mM) or Wnt 3a (200 ng/mL) or simultaneously with both treatment for 24 hours (*n*=4). Data are normalized and expressed as the % increase, or % of control, or fold of basal (x basal), or in relative amount in comparison to control, in mean ± SEM. ** *p* < 0.01.

**Supplementary Fig. 2.**
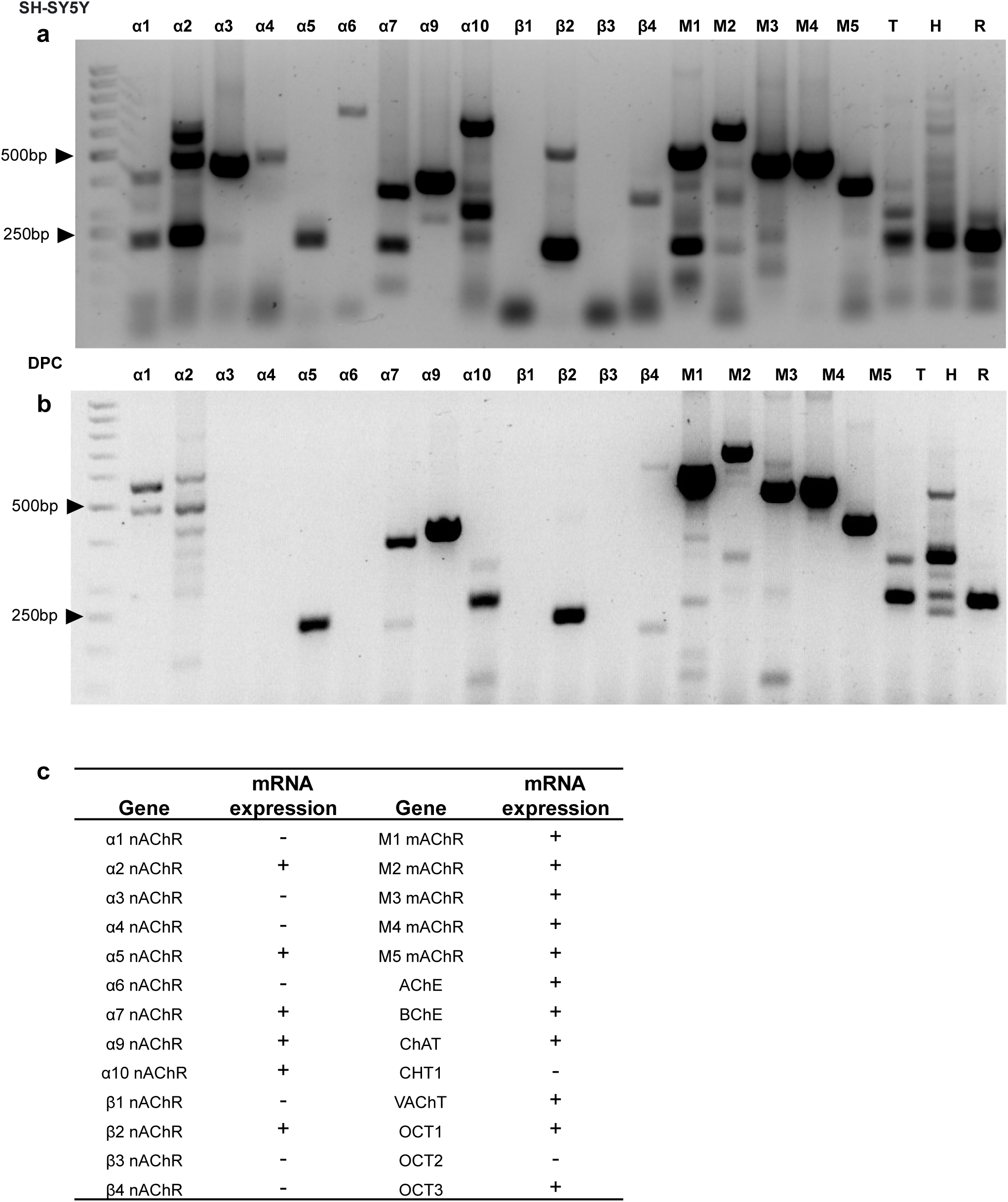
Expressions of mRNA encoding cholinergic molecules in DPC. **(a)** DNA gel electrophoresis of PCR product of various subunits of nicotinic and muscarinic receptor of AChR in SH-SY5Y (*n*=3). **(b)** DNA gel electrophoresis of PCR product of various subunits of nicotinic and muscarinic receptor of AChR in DPC (*n*=3). **(c)** Summary of mRNAs of cholinergic molecules in DPC (*n*=3).

**Supplementary Fig. 3.**
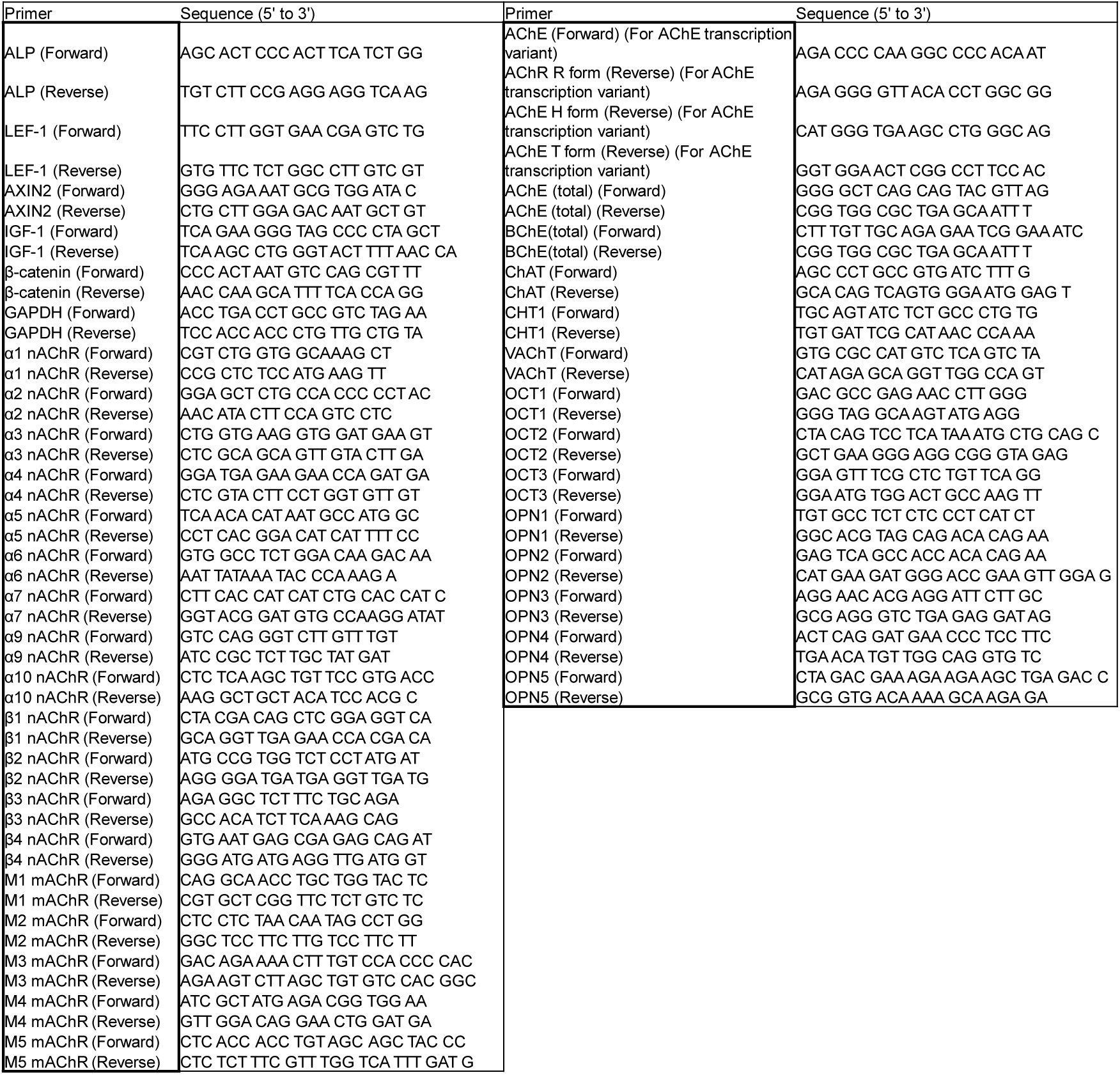
Sequences of primers for PCR. The sequences of primers are listed. nAChR: nicotinic acetylcholine receptor. mAChR; muscarinic acetylcholine receptor. AChE-R: readthrough acetylcholinesterase. AChE-H: hydrophobic acetylcholinesterase. AChE-T: tailed acetylcholinesterase. CHT1: high-affinity choline transporter-1. OCT: organic cation transporters. OPN; opsin.

